# CITEgeist: Accurate deconvolution of spatial transcriptomics with same-slide proteomics reveals midkine as a secreted microenvironment modulator in ESR1 mutant breast cancer

**DOI:** 10.1101/2025.02.15.638331

**Authors:** Alexander Chih-Chieh Chang, Brent T. Schlegel, Neil Carleton, Priscilla F. McAuliffe, Hunter Waltermire, Steffi Oesterreich, Russell Schwartz, Adrian V. Lee

## Abstract

**Background:** Dysplastic tissue architecture in estrogen receptor-positive (ER+) breast cancer across therapy-naïve and therapy-exposed cancer tissues presents unique challenges in the analysis of spatial transcriptomics. Many tools for deconvolution are developed on well-structured tissue architectures such as the 10x Genomics mouse hippocampus dataset. Spatial transcriptomics analysis could offer valuable insights into treatment response, but faces limitations in cellular resolution.

**Methods:** To address this problem, we developed CITEgeist, a computational tool for spatial transcriptomic deconvolution using integrated proteomics data from the same slide. Visium Antibody Capture technology was applied alongside our novel algorithm to analyze the tumor microenvironment. We demonstrate the reliability of our method using pre- and post-treatment samples from six breast cancer cases.

**Results:** Our approach revealed previously undetectable cellular interactions within the tumor microenvironment. By taking an interoperable approach to software development and grounding our algorithm in interpretable variables, we demonstrate how CITEgeist deconvolution is not only accurate but robust enough to be directly used as input in external analytical tools developed by other research teams. We then applied this approach to a set of specimens from a prospective trial our group ran and further validated the findings in a series of in vitro experiments as a demonstrated use case of the utility, necessity, and flexibility of CITEgeist; and the potential of our method to rapidly translate novel clinical samples to new biological insights.

**Conclusions:** CITEgeist addresses a critical technical gap in spatial multi-omics analysis through an integrated, multi-disciplinary approach. This work demonstrates the value of combining clinical, translational, and computational expertise to identify novel mechanisms of treatment resistance, potentially transforming therapeutic strategies for resistant disease.

## Background

Estrogen receptor-positive (ER+) breast cancer accounts for approximately 85% of new breast cancer diagnoses [1], with surgery and anti-estrogen therapy established as the first-line treatment [2]. How breast cancer changes post-treatment is of vital importance to disentangling how treatment resistance develops, and spatial transcriptomics offer a unique opportunity to dissect the mechanisms of cancer evolution in native tissue architectural contexts. Additionally, many techniques allow for the joint collection of proteomic information from the same slide, allowing integration of multi-omic informational landscapes, and hence, deeper insights.

However, no tool is currently capable of integrating this proteomic information for analysis, and there are also limitations in various single-cell reference-dependent deconvolution tools that have been the dominant design choice in the field.

Single-cell reference-based deconvolution methods have several disadvantages. First, it is costly to generate a single cell reference to match each spatial dataset, at approximately double the cost [3]. Second, external references or atlases are not always available, especially for heterogeneous post-treatment cancer samples, which in themselves are rare and difficult to collect routinely [4]. Third, biosamples such as core biopsies often provide too little tissue for single-cell sequencing [8], and, even if available, the biopsy tissue used for the reference may not capture the heterogeneity of the cancer slice used in the spatial transcriptomics. [5,6]. Fourth, there is a significant risk of data hallucination, where gene expression or cell-type abundance, as estimated from single-cell data, is artificially imposed in a spatial context, erroneously granting spatial properties to transcriptomic data when they are actually artifacts [7]. Finally, many methods suffer from a lack of interpretability, requiring complex assumptions and architectures to integrate the two informational landscapes of single-cell transcriptomics and spatial transcriptomics [8], which often is incompatible with other downstream analysis options and tools popular in the field, i.e., by failing to preserve raw counts needed in DeSeq [9].

These technical limitations prompted the development of CITEgeist, a novel computational tool that enables spatial transcriptomic deconvolution using integrated proteomics data obtained from identical tissue sections via integer linear programming. We demonstrate the accuracy of CITEgeist using a simulated reference test.

Additionally, we run this tool on post-treatment samples from a clinical trial examining endocrine therapy in older women who forego surgery to emphasize the rapid cutting-edge applicability of this method. Furthermore, we develop intrinsic tests on the non-simulated real tissue data to confirm the confidence of CITEgeist outputs on post-treatment tumor data. Lastly, we demonstrate how our output is fully compatible with tools developed by other groups, and use them to identify a novel environmental effect in ESR1 mutant breast cancer, which we then validate in cell culture in the laboratory.

## Methods

### CITEgeist Model

CITEgeist deconvolutes spatial transcriptomics data by leveraging antibody capture from the same tissue section, eliminating dependency on external single-cell RNA sequencing references. For the complete derivation, please see the Supplementary Methods. The method consists of three key components: (1) protein-based deconvolution using an Expectation-Maximization (EM) algorithm, (2) regional refinement with neighborhood optimization, and (3) spatial gene deconvolution with variable-sized neighborhoods.

### Software

CITEgeist is implemented in Python 3.10 using Gurobi 11.0.2 for Integer Linear Programming (ILP) optimization. Data preprocessing includes gene filtering (> 0 counts in ≥ 1% spots, mean expression > 1.1) and normalization (target sum 10,000). Antibody data undergoes winsorization for the top and bottom 5% of data points, followed by global centered log ratio (CLR) normalization. Complete mathematical derivations and implementation details are provided in Supplementary Methods.

### Single-Cell RNA-seq Reference

To establish a baseline for gene expression profiles — both for deconvolution tasks and the simulation of spatially resolved transcriptomic datasets — we utilized a publicly available atlas of single-cell transcriptomic data from human breast cancers, explicitly focusing on estrogen receptor-positive (ER+) tumors. This dataset served as the reference for deconvolution methods like Seurat [10] and cell2location [8], as well as the foundation for our simulations, enabling us to generate in-silico spatial datasets with a quantifiable ground truth of cell type proportions and gene expression levels.

We selected 12,000 cells from ER+ tumor samples across 11 ER+/HER2- cases in the Wu et al. breast cancer reference dataset [11], maintaining cell type proportions through random down-sampling from the primary atlas according to the annotations of 9 distinct cell types. Due to the computational constraints of the scCube autoencoder framework [12], we used this down-sampled version of the atlas as our primary reference for spatial transcriptomic simulations.

For the benchmarking of reference-based deconvolution methods, we utilized two distinct permutations of our reference, the first of which retained the complete set of ER+ tumor cells from the 11 patients from the Wu et al. atlas [11]. This ensured that the reference expression profiles for cell types were as comprehensive as possible, avoiding potential biases introduced by down-sampling. We then performed the same analysis using our second permutation of the reference, down-sampled from ∼30,000 to 8,000 cells to reflect both cancer heterogeneity and a realistic simulation of a single-sample scRNA-seq reference. By using both approaches, we ensured that our benchmarking fairly evaluated the deconvolution performance without artificial constraints on the reference profiles, while simultaneously reflecting realistic conditions in our simulated experiments.

### Spatial Transcriptomic Simulation

We utilized the scCube (v2.0.0) framework [12] to simulate Visium-like spatial transcriptomic data from the downsampled ER+ scRNA-seq atlas. A Variational Autoencoder (VAE) model with a hidden layer size of 128, a learning rate of 0.0001, and 10,000 epochs generated single-cell gene expression profiles for the nine major reference cell types. Cells were generated based on the proportions of the annotated cell types in the Wu atlas, with a batch size of 512, executed on a CUDA-enabled A100 GPU. The trained model was saved for future use.

To capture the spatial distribution of cells, each sample was modeled on a 50 × 50 spatial unit grid, divided into hexagonal Visium-like spots, with an average of five cells per spot. This structure enabled the modeling of clustered and infiltrative patterns typical of the breast tumor microenvironment. Clusters of cells representing various types were assigned specific spatial parameters such as shape, twist, and scale, while infiltration patterns were intentionally varied within and across clusters. Background cell types were also incorporated, with proportions reflecting approximate tumor heterogeneity.

### Spatial Antibody Capture Simulation

Spatial antibody capture data were simulated for each replicate by first defining two simulated protein markers for each unique cell type in the dataset, resulting in a set of cell-type-specific markers. For each simulated spatial spot, the proportion of each cell type was retrieved, and the expected expression for each marker was calculated by scaling the cell type proportion by a random factor drawn from a uniform distribution between 20 and 50. Expression values for each marker were then simulated using a negative binomial distribution with a mean proportional to the expected expression and a dispersion parameter of 0.5. A 5% dropout rate was applied to the simulated expression values, which set the expression to zero to simulate additional technical noise. Low-level background expression was drawn from a negative binomial distribution with lower-scale parameters for spots where a given cell type was absent. In addition to the cell type-specific markers, nonspecific proteins were simulated across all spots, drawing from a negative binomial distribution with a constant mean expression of 10. The final output was a matrix of simulated protein expression values for each marker across all spatial spots.

To comprehensively evaluate deconvolution methods, we simulated distinct sets of mixed and highly segmented populations, acknowledging that actual cancer data spans a continuum between these two states. By showcasing performance across these diverse tissue architectures, we offer a thorough assessment encompassing the full spectrum of tissue heterogeneity. Our analysis emphasizes both population types: the mixed populations, which are designed to more closely resemble complex tumor microenvironments, and highly segmented populations, which have been foundational in prior deconvolution method development.

In order to account for spatial variability, we generated five distinct spatial replicates for each sample type ("high_seg" and "mixed"), each exhibiting similar yet unique spatial patterns. This approach yielded spatial gene expression matrices at single-cell and spot-level resolutions, where each spot’s gene expression values reflect the aggregated profiles of the cells within it. Additionally, we recorded the cell-type proportions per spot to provide a ground truth for evaluating spatial deconvolution accuracy. These datasets, featuring diverse spatial configurations and infiltration patterns, aim to offer a more realistic foundation for assessing different deconvolution methods on tumor-like spatial data.

### Cell Culture

Establishment of MCF-7 and T47D mutant *ESR1* cell lines was previously reported [13]. Individual MCF-7 and T47D wild-type and CRISPR-edited D538G ESR1-mutant cell clones were evenly pooled and maintained in Dulbecco’s modified Eagle medium (ThermoFisher 11965118) or RPMI (ThermoFisher 11875119), respectively, plus 10% fetal bovine serum, and 1x penicillin-streptomycin (Sigma-Aldrich P4333) at 37°C in a humidified incubator with 5% CO2. For hormone treatment experiments, cells were deprived of steroid hormones by placement in phenol red-free Improved Minimum Essential Medium (Richter’s Mod.) (Fisher Scientific MT10026CV) plus 10% charcoal stripped serum (CSS). 17β-Estradiol (E2) (Sigma-Aldrich E2758-1G) was used at a final concentration of 1nM for +E2 conditions. ELISA was performed using the Invitrogen Human MDK ELISA Kit (EH319RB) according to manufacturer instructions.

### Immunofluorescence

MCF-7 and T47D cells were plated in 8-well chamber slides (FisherScientific 08-774-26) at 500 cells/well in full medium as described. After adhering to the wells for 24 hours, the medium was changed to phenol red-free Improved Minimum Essential Medium (Richter’s Mod.) (Fisher Scientific MT10026CV) plus 10% CSS for 5 days, with the medium changed 2 times per day. For +E2 conditions, 1nM E2 was added to the medium for 24 hours. The cells were fixed in 4% PFA (FisherScientific 50-980-487), permeabilized for 10 minutes in 0.2% Triton X-100 (Sigma-Aldrich X100-100ML) in PBS (Fisher Scientific MT21031CV), then blocked for 30 minutes in 1% BSA (Sigma-Aldrich A9647-500G), 10% goat serum (Sigma-Aldrich G9023), 0.1% Tween20 (Sigma P7949- 100ML) in PBS. Primary antibodies for anti-MDK (1:1,000 ThermoFisher MA5-32538) and anti-E-cadherin (1:200 BD Biosciences 610182) were diluted in 1% BSA, 0.1% Tween20 in PBS and incubated overnight at 4 C in a humidified chamber. The cells were then incubated in secondary antibodies for anti-rabbit Alexa Fluor 647 (Invitrogen A32733) and anti-mouse Alexa Fluor 555 (Invitrogen A32727) for 1 hour at room temperature in the dark. Counterstaining was performed with Hoechst 33342 (ThermoScientific 62249) in PBS for 10 minutes, and cells were imaged on a Nikon AX confocal microscope with 10 image z-stacks in a minimum of 3 locations per sample at 60x oil magnification. Quantification was performed with CellProfiler and ImageJ.

### Code Availability

The CITEgeist package, including the tool itself, benchmarking scripts, and code for simulating test datasets, is available on GitHub at https://github.com/leeoesterreich/CITEgeist. The repository provides the complete implementation for generating the spatially resolved CITE-seq data simulations and evaluating deconvolution accuracy and gene expression (GEX) profile inference performance. It includes workflows for benchmarking CITEgeist against state-of-the-art methods and instructions for replicating the analysis and customizing simulations for new datasets. Comprehensive documentation and example scripts are included to ensure usability and reproducibility. A persistent version of CITEgeist will be made available via FigShare at the time of publication (doi: 10.6084/m9.figshare.28385675).

### Clinical Trial Sample Collection

As part of a prospective, pragmatic, hybrid de-centralized non-randomized clinical trial (NCT05914792) [14] designed to evaluate circulating tumor DNA (ctDNA) levels as a means to augment longitudinal monitoring of older patients with ER+ breast cancer receiving primary endocrine therapy (pET), tumor tissue samples were collected at baseline and at the time of surgical intervention [15] . In brief, patients aged 70 years and older with ER+/HER2- non-metastatic breast cancer signed informed consent (protocol approved by the University of Pittsburgh IRB under STUDY21100091 and conducted through the UPMC Hillman Cancer Center under protocol 22-088) and chose to forego upfront surgery in favor of pET. Because this trial focused on surgical de-escalation, most patients did not undergo surgery during the study follow-up. However, a subset of six patients, four responding and two progressing, based on imaging assessment and ctDNA results, underwent surgery after 3-50 months of pET (12 total specimens). We collected matched core biopsy and surgical specimens from these six patients (12 total specimens) for this analysis, providing a use case for the CITEgeist tool. To preserve patient anonymity, all samples are labelled HCC22-088-P#-S#, for HillmanCancerCenter trial 22-088 -Patient#-Sample#. Sample number is chronological; in all cases, the biopsy samples are one, and the surgical samples are 2.

**For specific statistical and sequencing methods, please refer to our attached Supplementary Methods.**

## Results

### CITEgeist is uniquely suited for accurate deconvolution in mixed dysplastic cancer architectures

We developed CITEgeist, a computational method designed specifically for accurate cell-type deconvolution in heterogeneous cancer tissues, inspired by a unique opportunity from a clinical trial at our institution (Figure 1A). As illustrated in Figure 1B, our method integrates multiple data layers to achieve high-precision cell-type identification. The core algorithm implements a neighborhood-based approach that utilizes the same-slide proteomic and transcriptomic data to capture cellular context at local, regional, and global scales, providing a comprehensive view of the tumor microenvironment.

**Figure 1:**
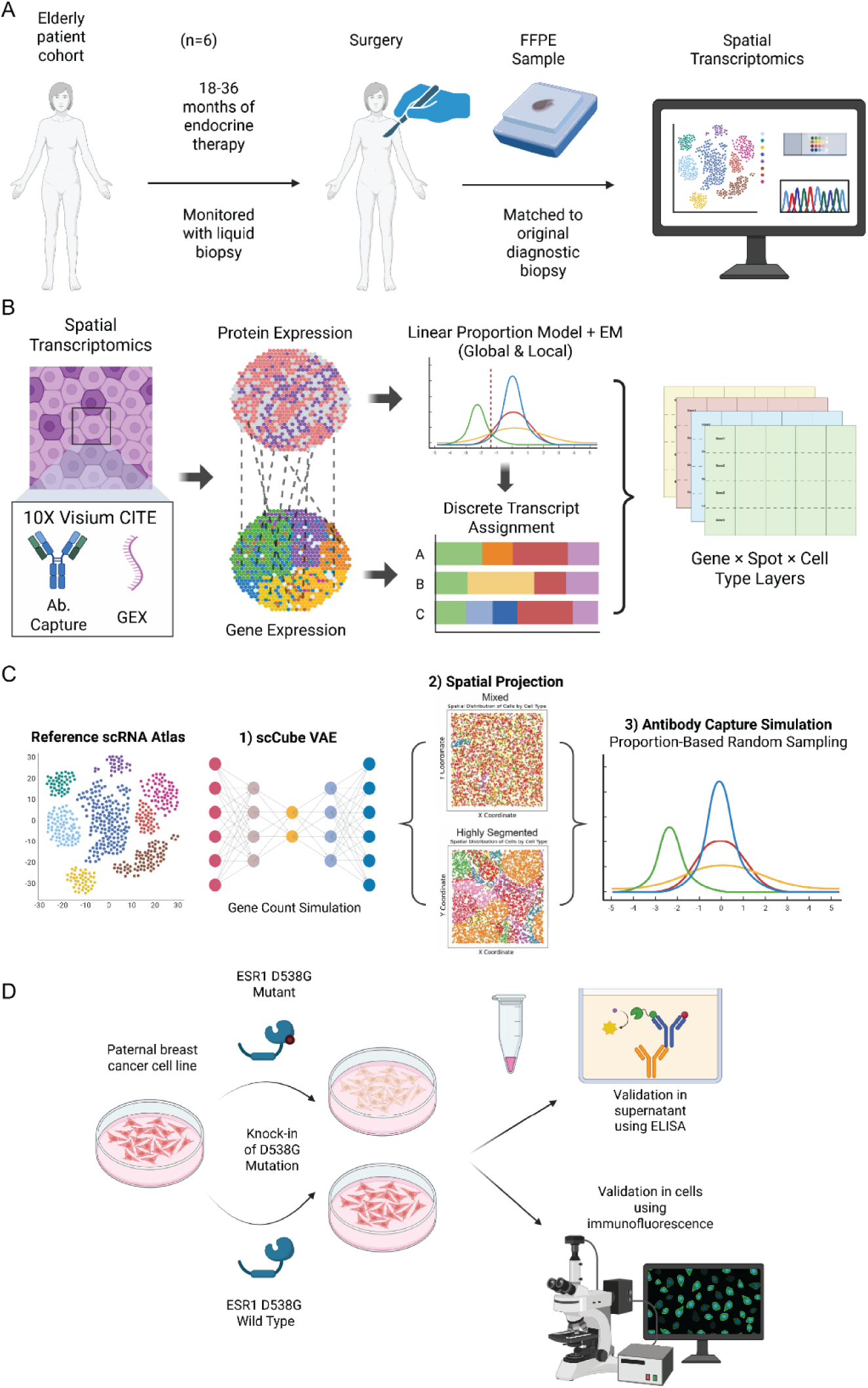
Graphical abstract of the CITEgeist development process. (A) Graphical abstract describing the clinical trial tissue collection process. (B) Graphical abstract describing the CITEgeist approach. (C) Graphical abstract describing the complete simulation pipeline for benchmarking CITEgeist against other state of the art single cell reference based spatial deconvolution methods. (D) Graphical abstract describing the wet lab validation process.

Unlike many existing tools, which are primarily validated on homogeneous tissues such as the mouse brain [8,16] we rigorously evaluated CITEgeist on cancer tissues with their characteristic heterogeneity. To this end, we generated a series of simulated datasets with varying degrees of cellular admixture and spatial segmentation (Figure 1C). These simulated benchmarks represent the complex architecture typical of dysplastic tissues, with intermingled malignant and non-malignant cells exhibiting different levels of differentiation. We then validate novel findings in cancer cell line models via ELISA and immunofluorescent imaging as outlined in Figure 1D.

### Benchmarking using simulated data demonstrates that CITEgeist outperforms state-of-the-art algorithms in deconvoluting cellular proportions

Using these simulated datasets, our comparative analysis revealed increased error in the mixed datasets compared to the segmented datasets with state-of-the-art methods such as Cell2Location and Seurat, both seeing increases in error from 0.08 RMSE to 0.167 RMSE and 0.10 RMSE to 0.133 RMSE for deconvoluting cellular proportions, respectively, when comparing highly segmented test sets vs mixed test sets. CITEgeist notably does not suffer from this drop, performing with a 0.08 RMSE in both circumstances.

To test the sensitivity of different methods to the size of the single cell reference, we also performed tests with reference datasets corresponding to single sample paired single cell sequencing (8,000 cells) or atlas-level objects (30,000 cells). Based on this test, RCTD was noted to have a severe reference dependency based on the comprehensiveness of the reference dataset used. All methods were first run with the comprehensive 30,000 cells, which in cost terms is about 4x the size of a standard single-cell sequencing run, and then run with 8,000 cells sampled from that distribution to capture what would be approximately as if a spatial sample had been submitted jointly with a singular single-cell sequencing kit. Most methods had minimal alterations in performance (Supplemental Files 2); however, RCTD showed deterioration under this test in both text contexts. With RMSEs of 0.21 on average with the 8,000 cell reference, compared to 0.05 with the 30,000 cell reference. This confirmed our hypothesis regarding the difficulty in evaluating performance on real cancer data when there are no clear guidelines on the necessary size and quality of single-cell references.

Tangram also notably and visibly proved a different issue of over-calling T-cells due to their known over-representation in single-cell datasets [17] (Figure 2B). It reported a JSD on average of 0.56 in the highly segmented dataset, and 0.53 in the mixed dataset, compared to an average of 0.15 and 0.34 in all other methods.

**Figure 2:**
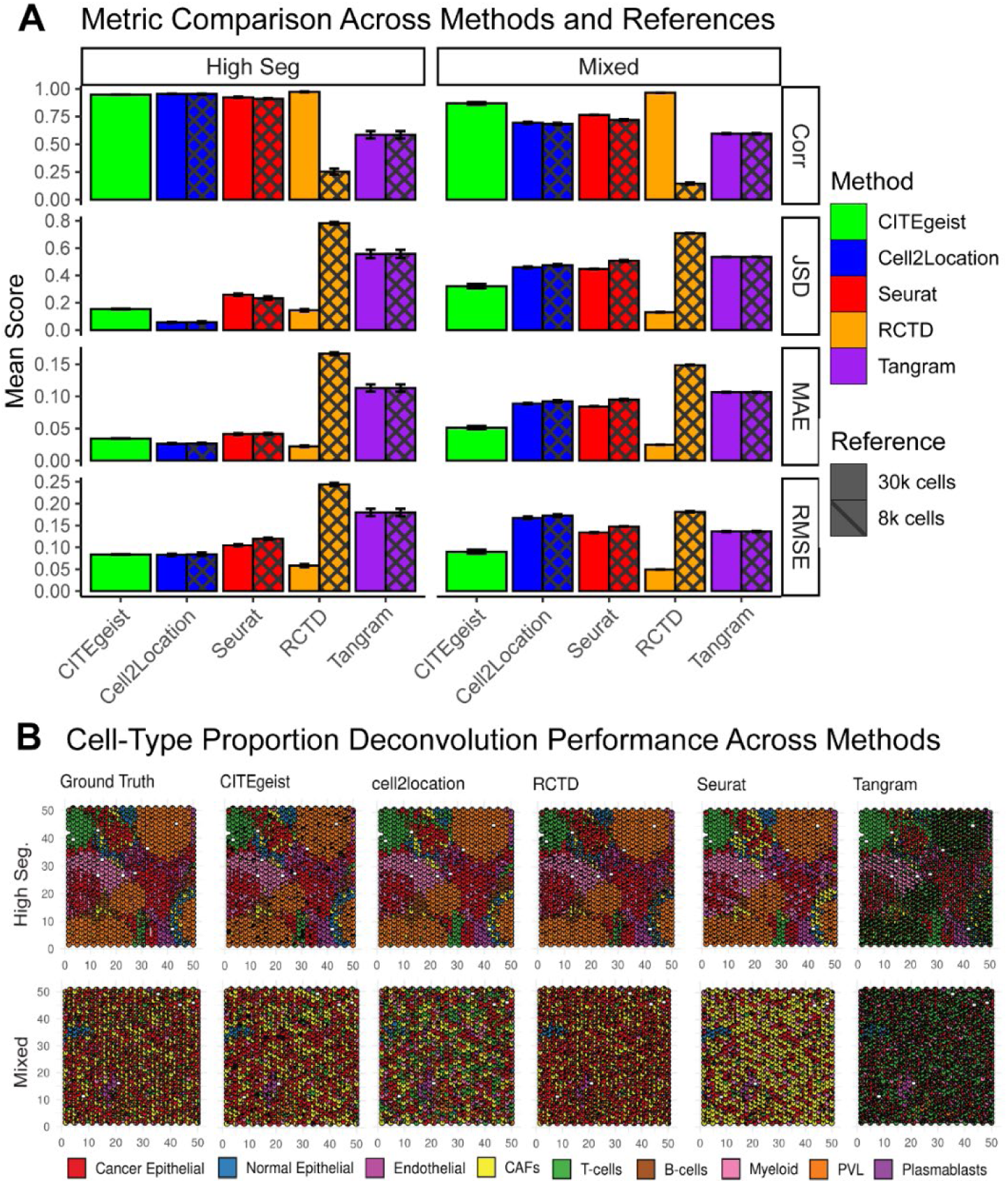
Performance evaluation of CITEgeist and reference-based cell type proportion deconvolution methods on simulated Visium datasets. (A) Quantitative comparison of deconvolution accuracy across different spatial patterns ("Highly Segmented" and "Mixed") and reference atlas sizes (30,000 or 8,000 cells). Metrics include Pearson’s Correlation (Corr, higher values indicate better performance), Jensen-Shannon Divergence (JSD), Mean Absolute Error (MAE), and Root Mean Square Error (RMSE), with lower values indicating better performance for the latter three error metrics. Error bars represent standard error (n = 5 independent replicates per condition). (B) Representative deconvolution results from all benchmarked methods showing cell-type proportion estimates across spatial spots for both Highly Segmented (top row) and Mixed (bottom row) tissue patterns. Each spot is colored according to the estimated proportions of nine cell types (legend at bottom). Ground truth (leftmost column) shows the actual cell-type distributions, while subsequent columns display predictions from each benchmarked method. Samples were selected from n = 5 independent simulations per condition to represent typical performance.

### CITEgeist deconvolutes gene expression counts and outperforms state-of-the-art algorithms

We then extended our benchmarking to evaluate the performance to deconvolute the spatially-derived gene expression of each cell type, a key function of spatial transcriptomics. At the time, only two methods had documented methods for deconvoluting RNA signals per cell type from their model, Cell2Location and Tangram (Figure 3).

**Figure 3:**
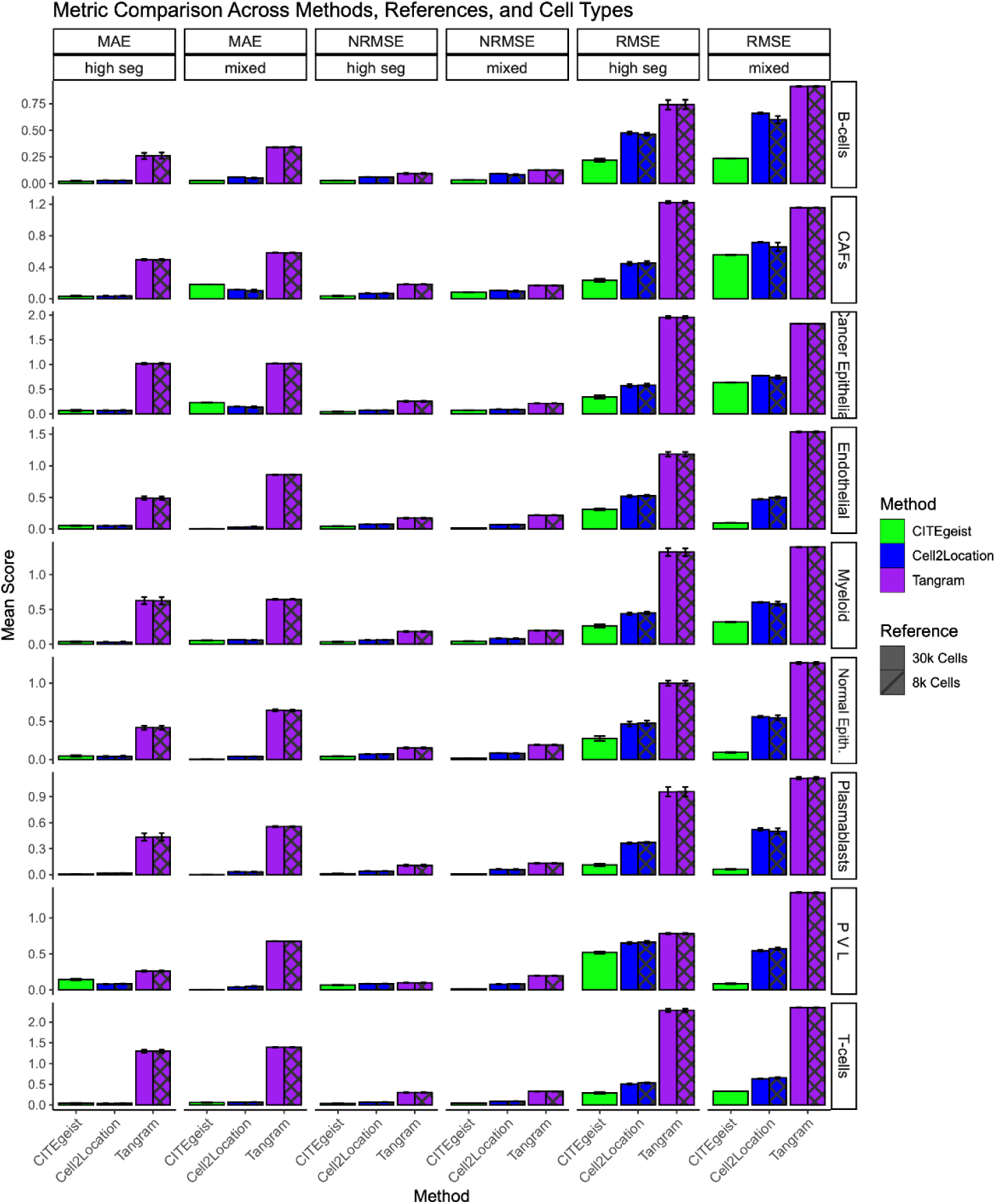
Metric comparison of gene expression (GEX) deconvolution per spot, evaluated across all benchmarked methods and reference cell types. Metric comparison of gene expression (GEX) deconvolution per spot, evaluated across all benchmarked methods and reference cell types. Metrics include Pearson’s Correlation (Corr, higher values indicate better performance), Jensen-Shannon Divergence (JSD), Mean Absolute Error (MAE), and Root Mean Square Error (RMSE), with lower values indicating better performance for the latter three error metrics. Error bars represent standard error (n = 5 independent replicates per condition).

In the mixed cancer-similar test set, CITEgeist had at least a 4-fold improvement over the other methods with an average NRMSE of 0.04, while Cell2Location had an average NRMSE of 0.16, and Tangram had an average NRMSE of 0.20 (Figure 3). We show a 2- to 4-fold increase in performance in the highly segmented test set (CITEgeist: 0.04 average NRMSE, Cell2Location: 0.066 average NRMSE, Tangram: 0.17 average NRMSE).

The magnitude of this significant performance improvement is visualized in Figure 3, which quantitatively illustrates CITEgeist’s accuracy metrics across different tissue composition scenarios compared to existing methods.

### Interpretability in CITEgeist output

A key issue in spatial deconvolution is also an over-reliance on simulated benchmarks [18]. Hence, interpretability and verifiable results are key, and thus we developed several intrinsically-validating tests for CITEgeist deconvoluted samples, as demonstrated in Figure 4, which allow for confidence in CITEgeist output on a sample-by-sample basis, independent of simulated testing or ground truth. Figure 4A illustrates our deconvoluted cell-type distributions, which were subsequently reviewed in conjunction with the underlying H&E image, demonstrating consistency in the results and expected proportions; with cancer cell proportion predictions matching prominent clusters of epithelial cells, fibroblasts matching fibrotic regions, and CD4 and CD8 T cells in expected proportions consistent with the known biology of these cell types.

**Figure 4:**
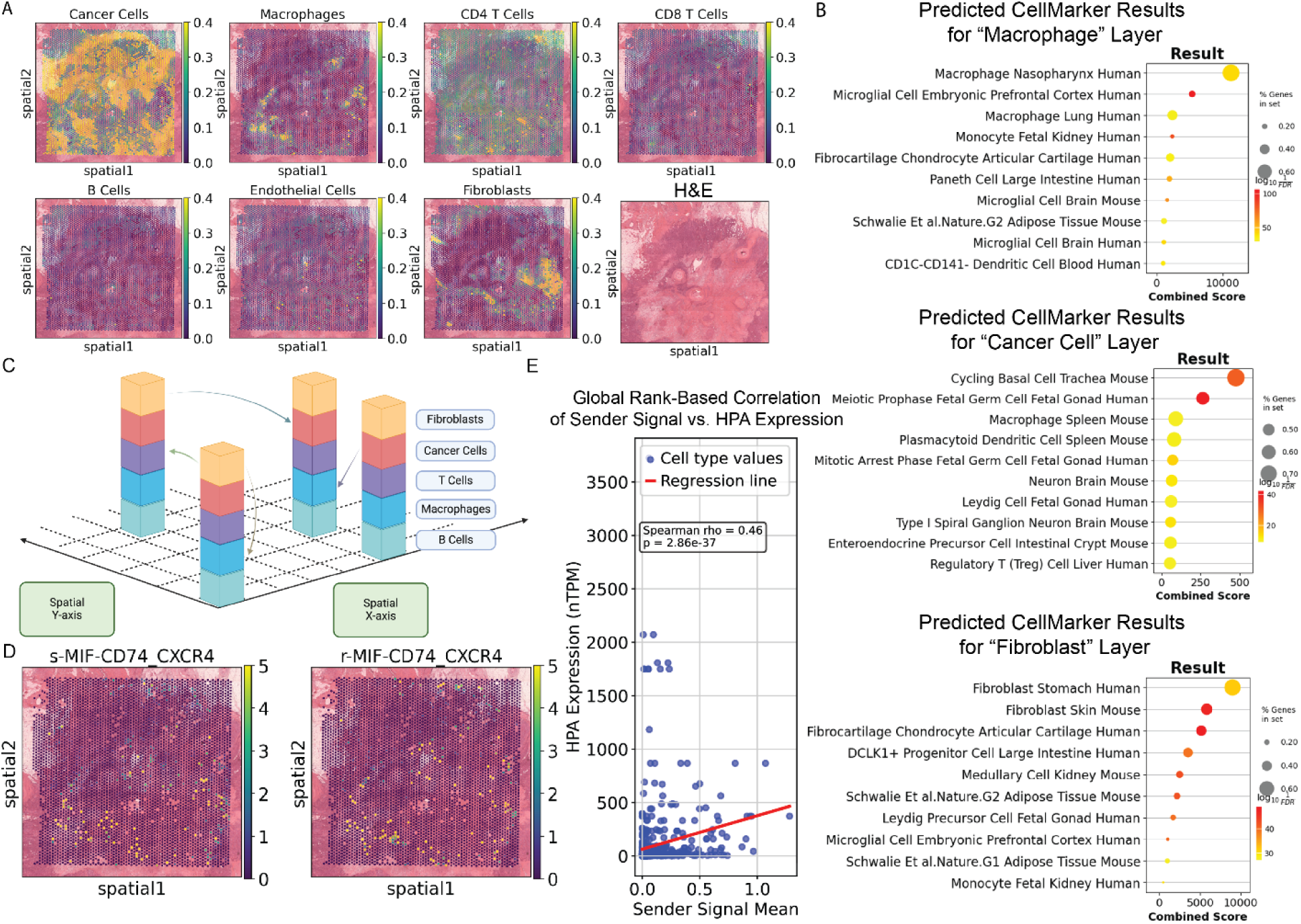
Demonstration of CITEgeist validity when applied to real cancer data. (A) Spatial plots showing the predicted proportions from CITEgeist output of the celltypes in sample HCC22-088-P1-S2 (B) CellMarker 2024 GSEApy calls, showing the top predicted celltypes for their respective ‘assigned’ gene expression layer. (C) Graphical abstract demonstrating how AnnData objects permit stacking of cell ‘compartments’ in the same adata spatial spot, and how COMMOT calculates using this information (D) Spatial plot showing an example of the detected sender and receiver signal for pathway MIF-CD74 CD44. (E). Global correlation regression plot showing the correlation between cell type assigned sender signal and said ligands expression in the Human Protein Atlas.

Additionally, to demonstrate the accuracy of the gene expression deconvolution, we took the gene layer for the three most abundant cell types: macrophages, cancer cells, and fibroblasts, and used the GSEApy package to assess against the CellMarker_2024 gene set, and showed excellent concurrence with 60%, 70%, and 60% gene set overlap respectively, as well as 3/3 aligned cell type calls for the top most pathway.

These methods demonstrate how even in the absence of ground truth, CITEgeist interpretability allows for excellent traceable results for downstream analysis.

### CITEgeist preservation of spatial context and RNA informational landscape allows for seamless integration with additional analysis tools

A key strength of CITEgeist is the interpretability of its output, facilitating seamless integration with various downstream analytical tools due to its use of linear assignment. Because we utilize strict integer assignments of RNA transcripts, we maintain what are effectively raw counts for programs such as DeSeq2 [19]. In Figure 4C, we illustrate the functional effect of CITEgeist deconvolution within the interoperable AnnData format. By stacking the deconvoluted compartments vertically and maintaining integer counts, we connect CITEgeist objects to COMMOT [19], a signaling prediction package designed for spatial data. We validated the biological relevance of CITEgeist results by examining all detected secretion signals and comparing them with established biological findings (Figure 4). To validate the accuracy of this output, we conducted a correlation analysis using the output from our samples for secreted signals and their assigned cell types, as well as known secretion amounts from the Human Protein Atlas (Figure 4E). This analysis showed a consistent correlation (Spearman Rho = 0.46, p-value = 2.86e-37).

### Signaling analysis using CITEgeist deconvoluted spatial transcriptomics identifies an upregulation in MDK signaling, confirmed with in-vitro experiments

After validating CITEgeist performance on simulated ground truth datasets and also real-world samples for interpretability, we applied CITEgeist to analyze clinical samples, focusing on a breast cancer case with an *ESR1 D538G* mutation initially identified through pilot bulk-RNA sequencing and subsequently confirmed by droplet digital PCR (ddPCR) (Supplementary Figure 1). This mutation was of particular interest, given the clinical relevance of *ESR1* mutations, and our laboratory’s previous research in this area^[13]^.

We have previously shown that *ESR1* mutant breast cancers express basal cytokeratins, and using CITEgeist on spatial transcriptomics from a case with an *ESR1* mutation, we confirmed the presence of basal genes in the Cancer Cell layer (Figure 5A). However, we were intrigued to note that these signals were not clustered in one location among the cancer cells as one would expect from a single subclone (Figure 5B), despite exhibiting all of the same gene mutation signatures from Estrogene [20] (Figure 5C) and known pathway alterations from our previous work (Figures 5D, 5E) [13]. As one of the first spatial samples of *ESR1* mutant breast cancer in the native breast microenvironment, we examined the signaling effect of these mutant breast cancer cells. We thus employed the validated COMMOT signaling analysis on CITEgeist-deconvoluted data, which revealed increased MDK, PTN, and MIF signaling in the mutant regions (Figure 5F). MDK has recently been shown to have a role in breast cancer, age, and tumorigenesis [21,22], and to validate our in silico findings, we performed ELISA on media from *ESR1* D538G MCF7 cell lines, and conducted analysis of the MDK immunofluorescence of MCF7 *ESR1* mutant cell lines. ELISA confirmed that the mutant cell line secreted double the MDK in environments both with and without estradiol (p < 0.0001) (Figure 5H). Additionally, the literature review identified that midkine has a strong pericellular effect, with a significant proportion being retained near the cell in non-cancerous settings [23]. Hence, we conducted image analysis of immunofluorescence on the same cell lines, which showed that in the vehicle, estrogen- deprived environment (akin to a treated patient), the mutant cell lines have approximately double the midkine at the cell membrane (p < 0.001) as well as in the cell as a whole (p < 0.01) (Figure 5I).

**Figure 5:**
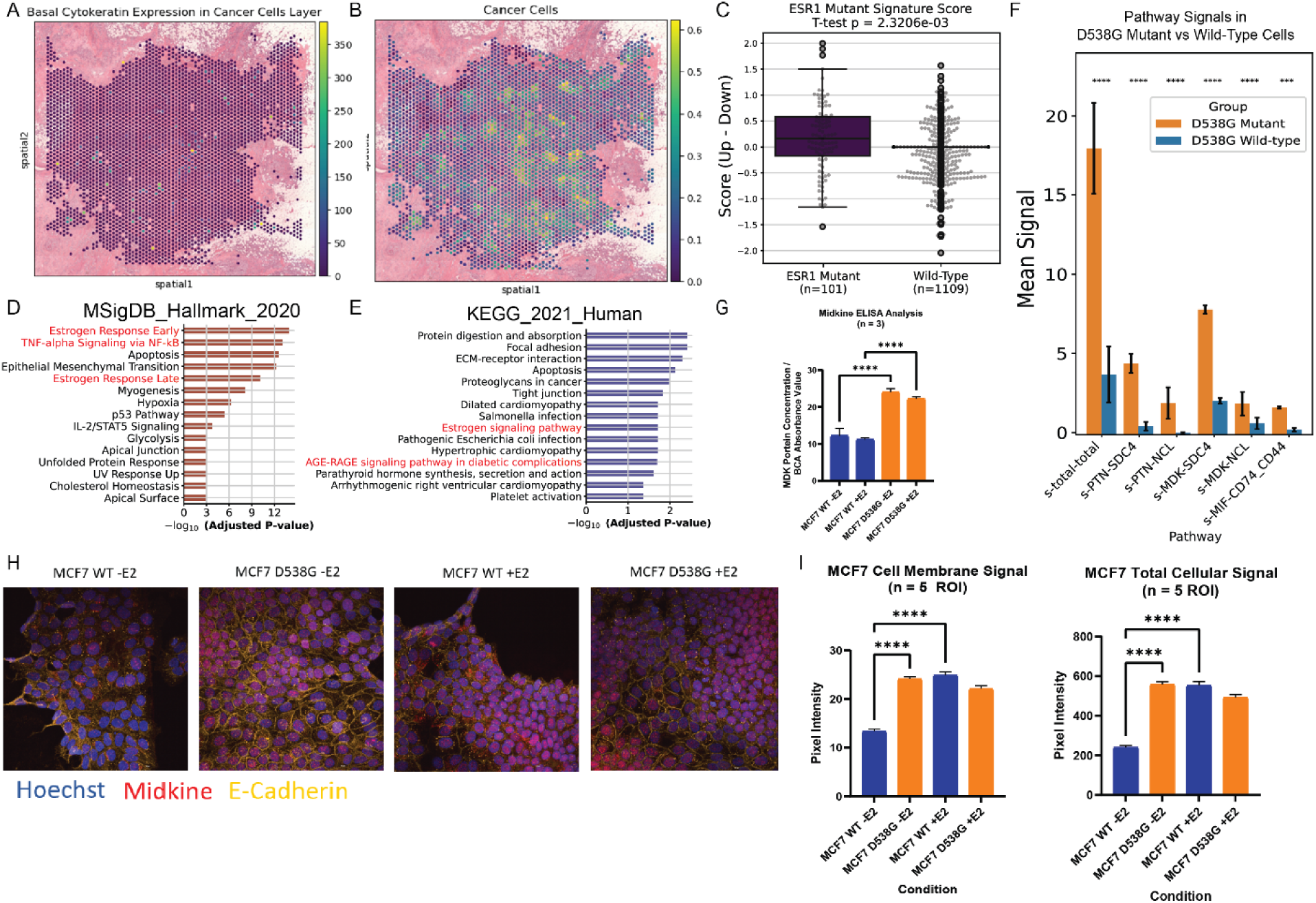
Wet lab validation of predictions based on CITEgeist output using external analytical packages. (A) Spatial plots showing the heterogenous deconvoluted result of elevated basal cytokeratin expression in the (B) Cancer Cells layer in the surgical sample from the same patient in sample HCC22-088-P4-S2, which has an ESR1 D538G mutation. (C) Combined box and scatter plot showing the average ESR1 Mutant Gene Signature Score between ESR1 D538G mutant cancer cells and WT cells. (T-test value 3.1229, p-value 2.3206e−03 (D) Gene pathways upregulated in the D538G cancer spots from the MSigDB Hallmark 2020 gene set. Red text pathways are previously confirmed upregulated activities in ESR1 mutant cells from Li et al. ^20^ (E) Gene pathways upregulated in the D538G cancer spots from the KEGG 2021 Human gene set. Red text pathways are previously confirmed upregulated activities from Li et al. (F) Bar plot showing the statistically significant COMMOT sender signal differences between D538G cancer spots and WT cancer spots. A Mann-Whitney U test was performed to compare pathway-specific sender signals between cells with and without the D538G mutation. Error bars are SEM. The two-way test statistic and p-value were computed, and pathways were ranked by significance. The mean difference in signal between mutation-positive and mutation-negative cells was calculated. Benjamini-Hochberg FDR correction was applied to control for multiple comparisons, and pathways with FDR < 0.05 were identified as significantly different. (G) Quantification of midkine ELISA results from MCF7 wild type and with ESR1 D538G mutation. Statistical testing was done with one-way ANOVA with Tukey’s post-hoc multiple comparison (H) Image examples of MCF7 WT and ESR1 D538G with and without 24 hours of E2 treatment after 5 days of E2 starvation. (I) Quantification of image examples assessing the midkine signal at the cell membrane and for the total cell signal, with values compared and tested via unpaired T-test. CellProfiler was used to assess values for each cell after Z Max Projection across 5 ROIs per slide.

## Discussion

Our study demonstrates that CITEgeist offers a robust computational approach for deconvoluting complex cellular architectures in cancer tissues. Here, we discuss the broader implications of our findings and contextualize them within the current landscape of spatial transcriptomics analysis.

### CITEgeist avoids many of the difficulties in using single-cell references in heterogeneous cancer environments

The integration of same-slide proteomics with our deconvolution method significantly enhances both interpretability and analytical robustness. By simultaneously capturing transcriptomic and proteomic profiles from the same tissue section, CITEgeist mitigates the technical variability inherent in comparing sequential slides, which is particularly valuable in heterogeneous cancer tissues where cellular composition can vary substantially across minimal spatial distances. This multi-modal approach provides internal validation and enables more confident cell-type assignment, especially in ambiguous cases where transcriptional profiles alone may be insufficient for accurate classification.

For example, we generated the reference size dependence test because Cell2Location has previously been demonstrated to be reasonably robust even in cases where the reference may not come from the exact same informational distribution [8], and hence wanted to replicate this test to test for the performance of each method in circumstances mimicking the effects of cancer heterogeneity. RCTD was noticeably challenged by this test, with a 4-fold increase in error – emphasizing the unique challenge of applying tools benchmarked on common public normal tissue datasets to a heterogeneous cancer dataset.

### CITEgeist is designed with interoperability and clinical considerations for maximum utility in analysis that allow for interpretable laboratory validation

A key contribution of our work lies in the methodology’s capacity to not only confirm existing biological knowledge but also to uncover novel insights during tool development. The identification of upregulated MDK signaling in *ESR1*-mutated breast cancer exemplifies how computational tools can drive biological discovery when designed with interpretability as a core principle. This finding emerged from our analytical pipeline rather than being a predetermined hypothesis, highlighting the value of data-driven approaches in uncovering unexpected biological mechanisms.

The *ESR1* D538G mutation identified in our study has significant implications for understanding tumor microenvironment dynamics. This mutation, occurring in the ligand-binding domain of the estrogen receptor, is known to confer constitutive activity and resistance to endocrine therapies. Our findings suggest that beyond its cell-autonomous effects, this mutation influences the broader tumor ecosystem through altered paracrine signaling, particularly via MDK. The increased MDK secretion observed in mutant cells potentially modulates immune cell recruitment, angiogenesis, and stromal remodeling, creating a microenvironment more conducive to tumor progression and therapy resistance. This expanded understanding of mutation effects beyond cancer cell-intrinsic properties represents an important shift in how we conceptualize the consequences of genomic alterations.

### Limitations

There are several limitations associated with CITEgeist. It is currently dependent on the variety and accuracy of the protein panel sequenced with spatial transcriptomics. It must be ordered together, which limits its utility in already sequenced datasets that lack protein information. However, as a first-in-class tool, it fills a necessary gap in the field where such samples already exist and have been sequenced, but researchers lack the tool to integrate proteomics and transcriptomics.

Secondly, our clinical trial has a small number of samples and is limited to breast cancer; hence, there may be other concerns or issues that emerge with other cancer types or tissue samples processed differently from other centers, and thus, careful usage and incorporation of feedback are critical to the long-term sustainability of the method.

Lastly, we also repeated our ELISA and imaging analysis in T47D, a different breast cancer cell line, which did not recapitulate the results (Supplemental Figure S3).

However, previous work has established that the effects of these mutants are often context-specific [24,25]. It is important to note that breast cancer is a notably heterogeneous disease [26], hence, the purpose of our wet lab validation is not to find something true for all breast cancer, but to demonstrate that CITEgeist output can be directly input into external tools developed by others, and still provide analysis that can be confirmed in an isogenic model, proving that CITEgeist deconvolution provides a robust statistical foundation for further abstractions via interoperable tool usage. Further work is needed to establish the whole pathway that supports increased midkine in certain ESR1 mutant breast tumors.

### Future Directions

Looking forward, several promising directions emerge from this work. First, the integration of temporal dynamics into spatial analysis would provide valuable insights into how signaling patterns evolve during disease progression and treatment response. Second, expanding our approach to incorporate additional data modalities, such as chromatin accessibility or metabolomic profiles, could further refine cell-type characterization and functional state assessment. Finally, the application of CITEgeist to larger patient cohorts across multiple cancer types would enable the identification of conserved and cancer-specific microenvironmental signatures, potentially revealing new therapeutic vulnerabilities.

Additionally, the computational framework established here could be adapted for other complex tissue architectures beyond cancer, such as in inflammatory conditions or developmental contexts, where cellular heterogeneity and spatial organization profoundly influence tissue function. The principles of neighborhood-based analysis and multi-modal integration are broadly applicable across diverse biological systems.

### Conclusions

In this study, we present CITEgeist, a computational method designed explicitly for accurate cell-type deconvolution in heterogeneous cancer tissues. Our approach demonstrates superior performance in mixed cellular architectures compared to existing tools, particularly in the challenging context of dysplastic cancer tissues. The integration of neighborhood context at multiple scales enables robust cell-type identification even in regions of high cellular admixture.

The interpretability of CITEgeist outputs facilitates seamless integration with complementary analytical tools, enhancing flexibility in downstream analyses. This interoperability allowed us to leverage CITEgeist’s deconvolution capabilities to identify and validate a novel biological finding: increased MDK signaling associated with the ESR1 D538G mutation, with potential implications for understanding endocrine resistance mechanisms.

By combining computational innovation with biological validation, our work contributes both a valuable methodological advancement and a specific mechanistic insight relevant to cancer biology. CITEgeist represents a step toward more comprehensive characterization of the complex cellular ecosystems in cancer, ultimately supporting the development of more effective therapeutic strategies that account for the full complexity of the tumor microenvironment.

### List of abbreviations

ESR1: Estrogen Receptor 1
CITEgeist: Cellular Indexing of Transcriptomes and Epitopes for Guided Exploration of Intrinsic Spatial Trends
MDK: Midkine

## Declarations

### Ethics approval and consent to participate

The protocol for the related clinical trial was approved by the University of Pittsburgh IRB under STUDY21100091 and conducted through the UPMC Hillman Cancer Center under protocol 22-088

### Consent for publication

Not applicable.

### Availability of data and materials

The Wu et al. breast cancer atlas can be accessed according to the instructions in their original publication; the processed scRNA-seq data utilized in this study can be downloaded through GEO (accession number: GSE176078, accessed 10/25/24). The downsampled reference and simulated test datasets will be available via FigShare at publication (doi: 10.6084/m9.figshare.28385675).

Visium datasets used in this paper will be published at the time of publication and are currently embargoed. Reviewers can access the data via the link and private access token provided to the journal in the cover letter at the time of submission.

All correspondence should be addressed to Adrian V. Lee at.

### Authors’ contributions

A.C.C. conceived the project, designed the computational framework, wrote the CITEgeist code, conducted the bioinformatic analyses, and prepared the initial manuscript draft. B.T.S. performed the CITEgeist benchmarking, simulation studies, and statistical analysis and contributed to discussing benchmarking and simulation strategies. N.C. facilitated the collection of clinical samples and coordinated the clinical trial. P.F.A. served as the clinical trial’s principal investigator and performed breast surgery, collecting the samples. H.W completed the wet lab validation of MDK signaling in ESR1 mutant cell lines. S.O. oversaw the overall project management and shaped the project’s goals. R.S. contributed to manuscript revisions and provided expertise in mathematical modeling and computational methodologies. A.V.L. guided the project design and objectives. All authors reviewed and edited the manuscript.

## Supporting information

Supplement 3 - Raw Data for Wet lab experiments

Supplement 1 - CITEgeist Proof

Supplement 2 - Benchmarking Data

## Acknowledgements

We want to thank Jingyang Qian for their gracious guidance in utilizing the scCube method to simulate and benchmark spatially resolved Visium-like datasets from a single-cell reference. We also thank Jian Chen from the Lee-Oesterreich Lab for her help with RNA sequencing prep and ddPCR.

We acknowledge the assistance of the physicians and clinical support staff at the University of Pittsburgh Medical Center health system for their contributions to the clinical trial—special thanks to Natera Inc. for their support with the clinical trial NCT05914792.

## Supplementary Files

Supplementary File 1: Full mathematical derivation of the CITEgeist method

Supplementary Files 2: Tables reporting the metrics of all compared and benchmarked methods.

Supplementary Files 3: Raw data from all wet lab experiments (ELISA and IF).

Supplementary Files 4: Supplementary Statistical and Sequencing Methods

## Supplementary Figures

**Figure S1:**
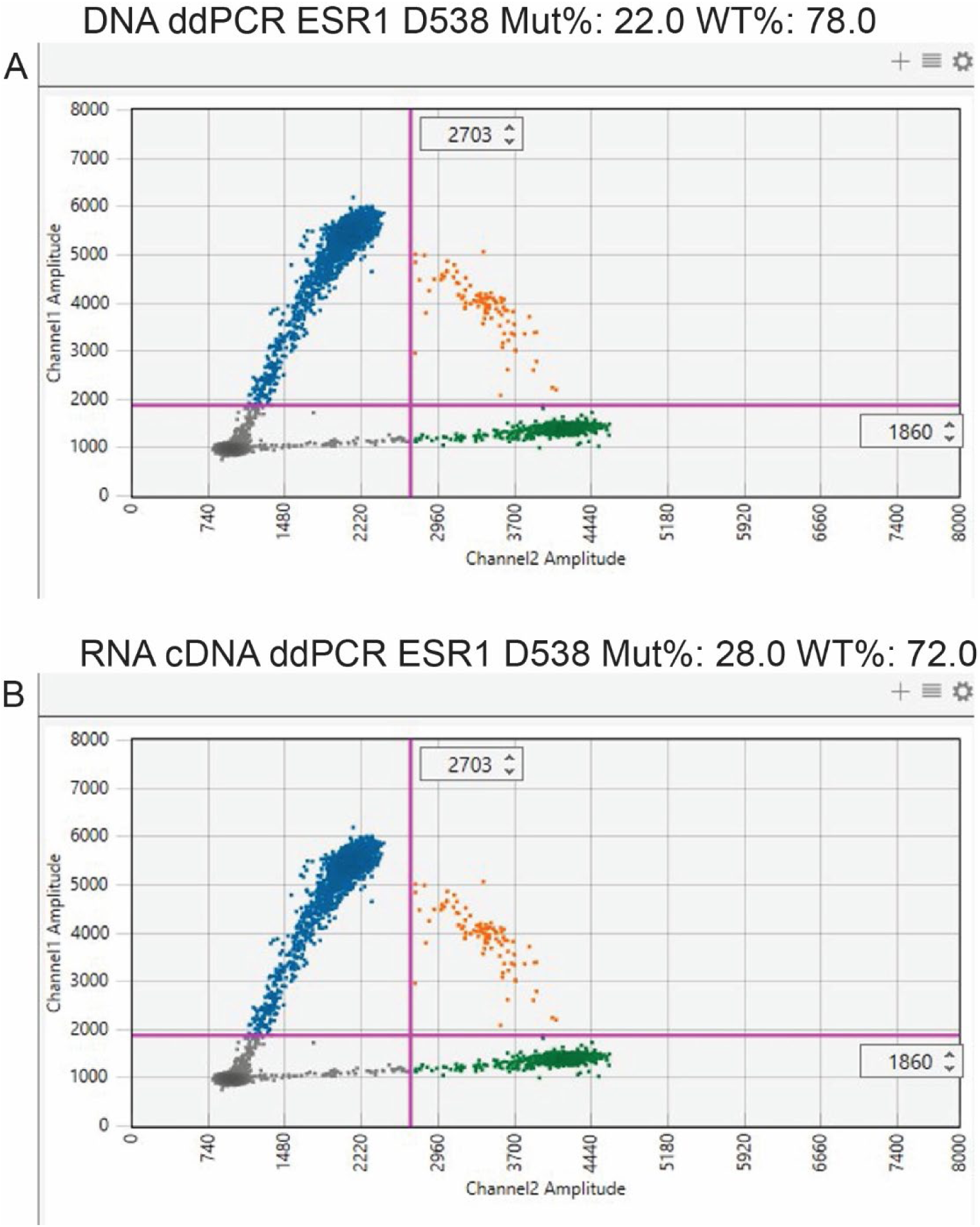
ddPCR confirming ESR1 mutation in HCC22-088-P4-S2. ddPCR confirmation of D538G mutation in Sample HCC22-088-P4-S2 (A) DNA, (B) RNA

**Figure S2:**
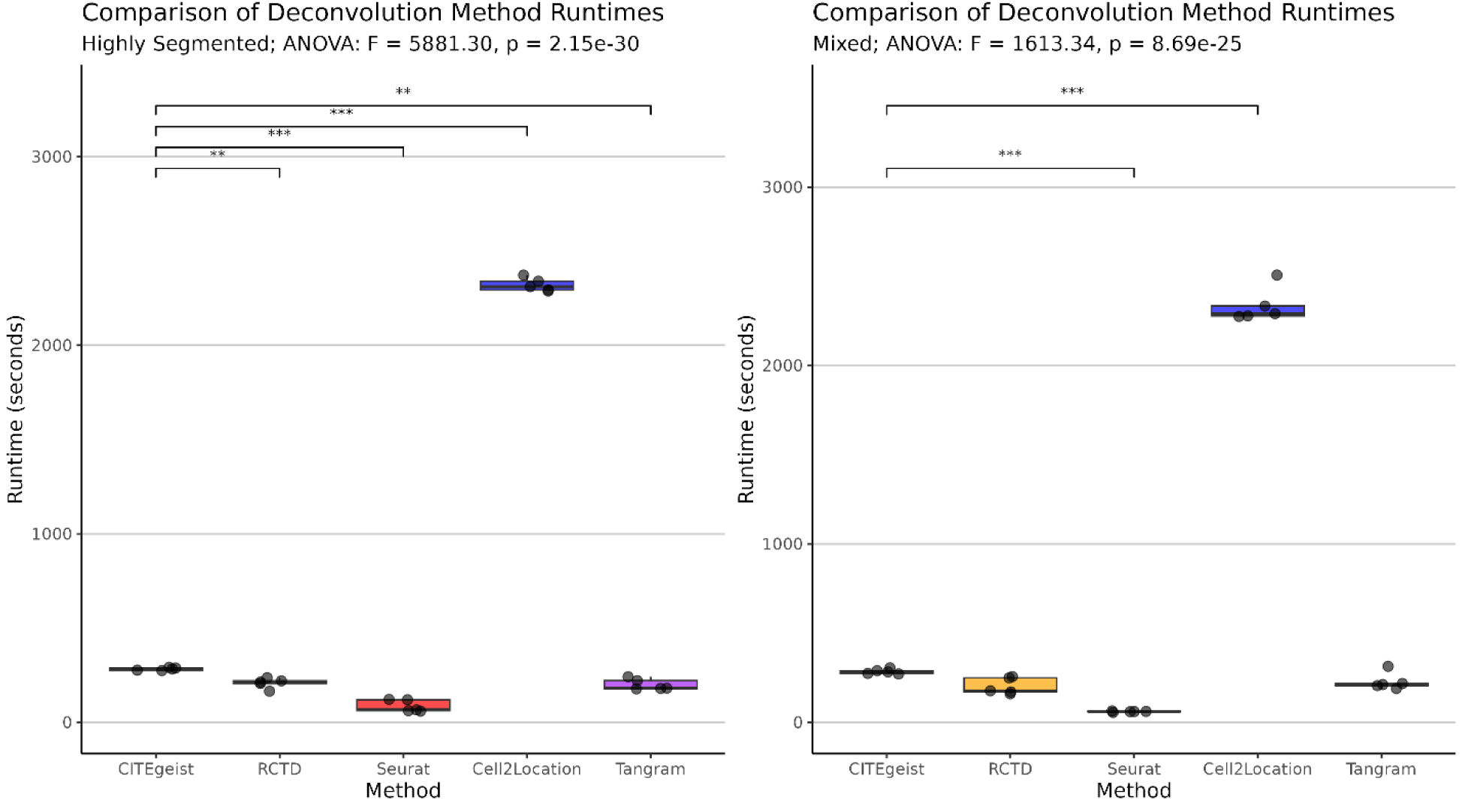
Runtime comparison of CITEgeist against other SOTA methods. Runtime comparison across all benchmarked methods, in both the “Highly Segmented” **(left)** and “Mixed” **(right)** simulated conditions. Statistical significance was calculated via ANOVA followed by Tukey’s Honest Significant Difference test.

**Figure S3:**
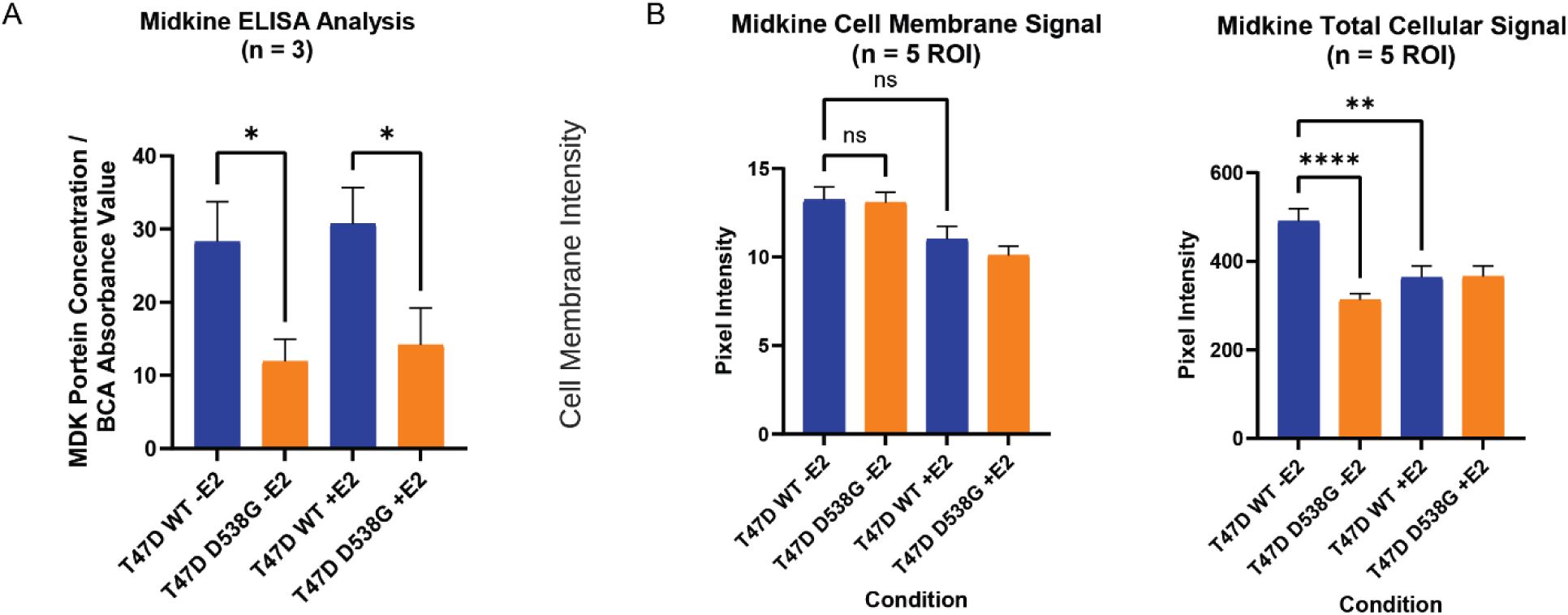
Quantification of midkine ELISA and immunofluorescence signals in T47D. (A) Quantification of midkine ELISA results from T47D wild type and with ESR1 D538G mutation. Statistical testing was done with one-way ANOVA with Tukey’s post-hoc multiple comparison. (B) Quantification of T47D midkine immunofluorescence assessing the midkine signal at the cell membrane and for the total cell signal, with values compared and tested via an unpaired T-test. CellProfiler was used to assess values for each cell after Z Max Projection.

