## Supplement 4 - Supplementary methods for "CITEgeist: Accurate deconvolution of spatial transcriptomics with same-slide proteomics reveals midkine as a secreted microenvironment modulator in ESR1 mutant breast cancer"

### Visium CytAssist v2 Library Preparation and Sequencing

Before sectioning, samples were assessed for RNA quality, and all samples had a DV200 score above the minimum threshold (≥30%). Spatial transcriptomic and proteomic libraries were generated with Visium CytAssist for the FFPE v2 gene expression kit (10x Genomics: 1000520) with the Human Immune Cell Profiling Panel (10x Genomics: PN-1000607).

Five-micron FFPE sections were placed on Schott Nexterion Hydrogel Coated Slides (Schott North America: 1800434) or Fisherbrand Superfrost Plus Slides (Fisher: 22-037-246) and processed through the 10x CytAssist for FFPE v2 protocol according to the manufacturer's instructions. Library QC was completed with an Agilent TapeStation 4150. Libraries were normalized and pooled to 2 nM before loading on an Illumina NextSeq 2000 using a P3 100 flow cell. The pooled library was loaded at 650 pM, and sequencing was carried out with a 28/10/10/50 base pair read structure, targeting 125 million reads per sample for transcriptomic libraries, and 25 million reads per sample for proteomic libraries.

### Total RNA Library Generation and Sequencing

RNA was extracted from FFPE sections using the Purelink FFPE RNA isolation kit (Invitrogen: K156002). According to the manufacturer's instructions, RNA-seq libraries were generated with the Takara SMART-Seq Stranded kit (Takara: 634447). RNA was normalized to 5 ng/µl in a total volume of 7 µl of input RNA. RNA fragmentation was not performed.

Ten cycles were used for PCR1, followed by depletion of ribosomal RNA using scZapR, and 12 cycles were completed for PCR2. Library quantification and evaluation were performed using a Qubit FLEX fluorometer and an Agilent TapeStation 4150. Libraries were normalized and pooled to 2 nM before sequencing on an Illumina NextSeq 2000 using a P4 200 flow cell. The pooled library was loaded at 750 pM. Sequencing was done with a 2×101 bp read structure, targeting 40 million reads per sample. A custom run chemistry was used to incorporate three dark cycles at the start of R2 to mask the three bases of the Takara adapter present in the read. Sequencing data was demultiplexed using the onboard Illumina DRAGEN FASTQ Generation software.

**ddPCR for D538G Mutation**

Droplet Digital PCR (ddPCR) was performed using the QX200 Droplet Digital PCR System (Bio-Rad) to detect the D538G mutation. Reactions were prepared using the ddPCR Supermix for Probes (No dUTP) (Bio-Rad, Cat. No. 1863024) in a duplex assay with a FAM-labeled probe targeting the mutant allele and a HEX-labeled probe for the wild-type allele.

Custom PrimeTime® qPCR probes and primers were synthesized by Integrated DNA Technologies (IDT) with HPLC purification. The wild-type allele was detected using a FAM-labeled probe with the sequence: /56-FAM/TC TAT GAC C/ZEN/T GCT GCT GGA GAT GCT /3IABkFQ/, while the mutant allele was detected using a HEX-labeled probe with the sequence: /5HEX/TC TAT GGC C/ZEN/T GCT GCT GGA GAT GCT /3IABkFQ/.

Droplet generation was performed using the QX200 Droplet Generator (Bio-Rad). Following PCR, droplets were read using the QX200 Droplet Reader, and fluorescence signal intensities were analyzed using QX Manager Software (Standard Edition). The FAM to HEX fluorescence ratio was used to determine the proportion of mutant and wild-type alleles in each sample.

Reagents used in this assay included the ddPCR Supermix for Probes (No dUTP) (Bio-Rad, Cat. No. 1863024) and custom PrimeTime® qPCR probes synthesized by IDT.

**Statistics for correlating Human Protein Atlas with CITEgeist deconvoluted signaling results**

Sender cell signals were extracted from a preprocessed AnnData object and assigned to the corresponding HPA gene expression values through a predefined dictionary that aligns cell types (e.g., 'CD8 T cells' and 'CD4 T cells' to 'T cells'). For each pathway—after removing irrelevant entries and extracting the ligand component for each pathway, we computed the mean sender signal for each mapped cell type. We obtained the corresponding average HPA nTPM value. Pathways with fewer than two matching cell types were excluded. For the remainder, Spearman rank correlations were calculated to assess the monotonic relationship between sender signal intensities and HPA expression. Finally, sender signal and HPA values were aggregated across pathways to compute a global Spearman correlation, visualized with a scatter plot overlaid by a linear regression line.

**ESR1 Mutant Signature Validation**

Gene lists corresponding to up- and down-regulated ESR1 mutant-associated genes were first extracted from the Estrogene 2.0 supplemental files [12] and filtered to retain the 20 genes in the cancer spatial dataset. For each spatial spot, an ESR1 signature score was computed as the difference between the mean expression of the upregulated genes and that of the down-regulated genes, with the resulting score appended to the metadata. Spots were then classified as ESR1 mutant or wild-type based on the "D538G Mutation" annotation, and the signature scores between these groups were compared using an unequal variance t-test, with 95% confidence intervals estimated via the t-distribution. Finally, the distribution of scores across groups was visualized using combined box and swarm plots.

**External Tools**

COMMOT [16], GSEAPY [17], and PyDeSeq2 [18] were used as described in their respective vignettes. COMMOT was run with a 1% cutoff for signals screened. GSEAPY gene lists were derived from Scanpy [19] Wilcoxon Rank Gene Group results with an adjusted p-value less than 0.05. PyDeSeq2 was run with size factors fit type set to 'poscounts' due to the sparse nature of spatial data.

**Benchmarking to State of the Art Reference-Based Deconvolution**

We benchmarked CITEgeist against several state-of-the-art tools for spot-wise cell type proportion estimation and deconvolution, namely: Seurat, RCTD, cell2location, and Tangram. These selected methods span the diversity of reference-based approaches to deconvolution, from RCTD's probabilistic framework to Tangram's deep learning approach via variational autoencoders. Notably, RCTD and cell2location have been lauded in previous benchmarking studies for their reported accuracy and stability in estimating ground-truth cell type proportions from Visium data.

Each method was implemented following the standard pipeline procedures detailed in their respective methodological vignettes, and all associated scripts for each method are available via the CITEgeist GitHub repository.

**Evaluation and Benchmarking**

We calculated several complementary metrics to evaluate the performance of the gene count prediction and cell type proportion models. These metrics provide a comprehensive assessment of the model's ability to predict gene counts and cell type compositions accurately.

**Evaluation of the Prediction of Gene Counts**

For the gene count predictions, we computed the root mean squared error (RMSE), normalized root mean squared error (NRMSE) and mean absolute error (MAE) between the ground truth and predicted gene counts. Raw gene count matrices were log(1 + x) normalized before metric calculation. The NRMSE normalizes the RMSE to the range or mean of the ground truth counts, allowing for comparison across cell types with different gene expression magnitudes. We calculated these metrics and reported the average and median RMSE, NRMSE, and MAE across all cell types.

**Evaluation of the Prediction of Cell Type Proportions**

For the cell type proportion predictions, we calculated the root mean squared error (RMSE) between the true and predicted cell type proportions, as well as the Jensen-

Shannon Divergence (JSD), mean absolute error (MAE), and Pearson correlation coefficient.

We also measured the JSD between the true and predicted cell-type composition vectors for each spatial spot. We reported the median JSD for all spots as the primary metric.

Additionally, we calculated the Pearson correlation coefficient between the true and predicted cell type proportions to evaluate the linear relationship between the two.

**Significance Testing**

To evaluate differences in performance across various deconvolution methods for spot-level cell type deconvolution, we performed a one-way analysis of variance (ANOVA). The method was the independent variable, while the selected metric was the dependent variable. Post-hoc pairwise comparisons were conducted using Tukey's Honest Significant Difference (HSD) test to identify specific group differences. Similarly, we utilized one-way ANOVA and post-hoc Tukey's HSD to determine significant differences in metrics between cell2location, Tangram, and CITEgeist for the GEX layer deconvolution task. Statistical significance was determined at $p<0.05$, with adjusted p-values reported for multiple comparisons.
