## Supplement 1 - CITEgeist Proof for "CITEgeist: Accurate deconvolution of spatial transcriptomics with same-slide proteomics reveals midkine as a secreted microenvironment modulator in ESR1 mutant breast cancer"

### Supplementary Methods

#### Detailed Mathematical Framework of CITEgeist

##### Data Preprocessing and Normalization

Prior to model application, we implement the following preprocessing steps:

1. Gene Expression Filtering:
   - Filter genes to those with count $> 0$ in $\geq 1.0\%$ of spots
   - Require mean expression $> 1.1$ in nonzero spots
   - Filter spots to minimum UMI count $\geq 100$
   - Normalize counts to target sum of 10,000
2. Antibody Data Processing:
   - Winsorize to remove top and bottom 5% outliers
   - Apply global centered log ratio normalization

##### Detailed EM Algorithm for Protein Deconvolution

The complete derivation of our EM algorithm follows these steps:

**Initialization** Let $N$ be the number of spatial spots, $T$ the number of cell types, and $S \in \mathbb{R}^{N \times T}$ the antibody capture matrix. Initialize:

- $Y^{(0)} \in \mathbb{R}^{N \times T}$ with random values between 0 and 1
- $\beta^{(0)} \in \mathbb{R}^{T}$ with all ones

**E-step Derivation** The complete objective function for the E-step includes:

$$\begin{aligned} L(Y|\beta) = &\sum_{i=1}^{N}\sum_{j=1}^{T}(S_{ij} - \beta_j Y_{ij})^2 + \\ &\lambda(\alpha \sum_{i=1}^{N}\sum_{j=1}^{T}|Y_{ij}| + (1-\alpha)\sum_{i=1}^{N}\sum_{j=1}^{T} Y_{ij}^2) \end{aligned} \tag{1}$$

Subject to constraints:

$$0 \leq Y_{ij} \leq 1 \quad \forall i, j$$

$$0.9 \leq \sum_{j=1}^{T} Y_{ij} \leq 1.2 \quad \forall i$$

**M-step Derivation** For the M-step, we derive the optimal $\beta_j$ by taking the derivative of the objective function with respect to each $\beta_j$ and setting it to zero:

$$\frac{\partial}{\partial \beta_j} \sum_{i=1}^{N} (S_{ij} - \beta_j Y_{ij})^2 = 0 \tag{2}$$

Leading to:

$$\beta_j = \frac{\sum_{i=1}^{N} S_{ij} Y_{ij}}{\sum_{i=1}^{N} Y_{ij}^2}$$

**Neighborhood Optimization Details**

**Neighborhood Definition** For each spot $i$, define neighborhood $\mathcal{N}(i)$ as:

$$\mathcal{N}(i) = \{j : d(i,j) \leq r\}$$

where $d(i,j)$ is the Euclidean distance between spots $i$ and $j$, and $r$ is the user-defined radius.

**Iterative Update Constraints** The constraint parameter $\delta = 0.4$ ensures stable updates:

$$\max(0, Y_{ij}^{(t)} - 0.4) \leq Y_{ij}^{(t+1)} \leq \min(1, Y_{ij}^{(t)} + 0.4)$$

**Local $\beta$ Updates** For each neighborhood:

$$\beta_j^{\text{new}} = \frac{\sum_{k \in \mathcal{N}(i)} S_{kj} Y_{kj}}{\sum_{k \in \mathcal{N}(i)} Y_{kj}^2}$$

##### 0.0.1 Complete Gene Expression Deconvolution Framework

**Expression-Aware Enrichment Score** For each gene $k$:

1. Define expression vector $\mathbf{g}_k$
2. Calculate threshold $\theta_k$:

$$\theta_k = \begin{cases} \text{median}(\mathbf{g}_k \mid \mathbf{g}_k > 0), & \text{if } \exists i : g_{ki} > 0 \\ 0, & \text{otherwise} \end{cases}$$

3. Define high-expression set $H_k$:

$$H_k = \{i \mid g_{ki} \geq \theta_k\}$$

4. Calculate normalized proportions:

$$\tilde{P}_{it} = \frac{P_{it}}{f_t + \epsilon}$$

5. Compute means:

$$\bar{P}_t^{(H)} = \frac{1}{|H_k|} \sum_{i \in H_k} \tilde{P}_{it}$$

$$\bar{P}_t^{(\text{bg})} = \frac{1}{N} \sum_{i=1}^{N} \tilde{P}_{it}$$

6. Calculate final enrichment:

$$E_t = \frac{\bar{P}_t^{(H)}}{\bar{P}_t^{(\text{bg})} + \epsilon}$$

$$E_t^{(\text{smooth})} = 0.8E_t + 0.2$$

$$E_t^{(\text{final})} = \frac{E_t^{(\text{smooth})}}{\sum_{t=1}^{T} E_t^{(\text{smooth})} + \epsilon}$$

**Complete ILP Formulation** The complete integer linear program for gene count allocation:

$$\begin{aligned}
\text{maximize} \quad & \sum_{j,m} (\text{gene_specific_enrichment}_{m,j} \times \\
& \text{cell-type_preference}_j \times \\
& \text{normalized_weights}_j \times \\
& \text{tie-break factor}) \times X_{j,m} \\
\text{subject to} \quad & \sum_{j=1}^{T} X_{j,m} = \text{center_counts}(m) \\
& X_{j,m} \geq 0, \ \text{integer} \\
& \text{tie-break factor} \sim U[0.9, 1.1]
\end{aligned}$$

**Optional Prior Integration** When using prior matrix $\mathbf{P}^{(\text{prior})}$, add penalty term:

$$-\lambda_{\text{prior\_weight}}(1 - P_{j,m}^{(\text{prior})}) \times X_{j,m}$$
